# Temperature increases soil respiration across ecosystem types and soil development, but soil properties determine the magnitude of this effect

**DOI:** 10.1101/2020.10.06.327973

**Authors:** Marina Dacal, Manuel Delgado-Baquerizo, Jesús Barquero, Asmeret Asefaw Berhe, Antonio Gallardo, Fernando T. Maestre, Pablo García-Palacios

**Affiliations:** Departamento de Biología y Geología, Física y Química Inorgánica, Universidad Rey Juan Carlos, C/ Tulipán s/n, 28933 Móstoles, Spain; Instituto Multidisciplinar para el Estudio del Medio “Ramon Margalef”, Universidad de Alicante, Carretera de San Vicente del Raspeig s/n, 03690 San Vicente del Raspeig, Spain; Departamento de Sistemas Físicos, Químicos y Naturales, Universidad Pablo Olavide, 41704 Sevilla, Spain; Department of Life and Environmental Sciences; University of California, Merced CA 95343, USA; Departamento de Ecología, Universidad de Alicante, Carretera de San Vicente del Raspeig s/n, 03690 San Vicente del Raspeig, Spain; Instituto de Ciencias Agrarias, Consejo Superior de Investigaciones Científicas, Serrano 115 bis, 28006, Madrid, Spain

**Keywords:** climate warming, land carbon-climate feedback, microbial biomass, nutrient availability, soil chronosequences, soil texture

## Abstract

Soil carbon losses to the atmosphere, via soil heterotrophic respiration, are expected to increase in response to global warming, resulting in a positive carbon-climate feedback. Despite the well-known suite of abiotic and biotic factors controlling soil respiration, much less is known about how the magnitude of soil respiration responses to temperature changes over soil development and across contrasting soil properties. Here, we investigated the role of soil development stage and soil properties in driving the responses of soil heterotrophic respiration to increasing temperatures. We incubated soils from eight chronosequences ranging in soil age from hundreds to million years, and encompassing a wide range of vegetation types, climatic conditions, and chronosequences origins, at three assay temperatures (5, 15 and 25°C). We found a consistent positive effect of assay temperature on soil respiration rates across the eight chronosequences evaluated. However, soil properties such as organic carbon concentration, texture, pH, phosphorus content, and microbial biomass determined the magnitude of temperature effects on soil respiration. Finally, we observed a positive effect of soil development stage on soil respiration that did not alter the magnitude of assay temperature effects. Our work reveals that key soil properties alter the magnitude of the positive effect of temperature on soil respiration found across ecosystem types and soil development stages. This information is essential to better understand the magnitude of the carbon-climate feedback, and thus to establish accurate greenhouse gas emission targets.

## Introduction

Temperature is a key driver of heterotrophic soil respiration (hereafter soil respiration), –a major process of carbon (C) loss to the atmosphere (Bond-Lamberty, Bailey, Chen, Gough, & Vargas, 2018; Bond-Lamberty & Thomson, 2010; Zhou et al., 2016). Global warming is expected to accelerate the rate of soil respiration (Davidson & Janssens, 2006; Kirschbaum, 2006), reinforcing climate change with a land C-climate feedback embedded in the Intergovernmental Panel on Climate Change (IPCC) projections (Ciais et al., 2014). Despite the recognized importance of an accurate representation of this feedback in Earth System Models to establish appropriate greenhouse gas emission targets (Bradford et al., 2016), the extent to which climate change will increase soil C losses to the atmosphere via soil respiration is still highly uncertain (Arora et al., 2013; Exbrayat, Pitman, & Abramowitz, 2014). Learning more about how and why soil properties regulate the magnitude of soil respiration responses to elevated temperatures is essential to accurately predict the land C-climate feedback in a warmer world.

To build confidence in the projected magnitude of the land C-climate feedback, the response of soil respiration to climate warming should be addressed across large spatial scales and encompassing a wide range of soil development stages. Beyond temperature, it is also critical to determine the influence of other key abiotic and biotic factors that regulate soil respiration (Guo et al., 2017; Rustad, Huntington, & Boone, 2000; Schindlbacher, Schnecker, Takriti, Borken, & Wanek, 2015). These include key soil abiotic drivers such as organic carbon (SOC), texture (i.e., the percentage of sand, silt, and clay), pH, and phosphorus (P), as well as biotic properties such as microbial biomass (Bradford, Watts, & Davies, 2010; Karhu et al., 2014). For instance, soil texture influences soil respiration by controlling water and nutrient availability (Delgado-Baquerizo et al., 2013) and regulating the potential of soil minerals to physically and chemically stabilize organic carbon (Rasmussen et al., 2018). A previous study showed that soils with higher proportion of clay sized particles also had higher microbial activity due to greater water and nutrient availability, leading to higher soil respiration (Karhu et al., 2014). Further, soil respiration increases as microbial biomass rises (Wang, Dalal, Moody, & Smith, 2003). Despite the knowledge accumulated about soil respiration drivers, much less is known about how soil properties modulate soil respiration responses to warming.

Soils are known to develop from centuries to millennia, resulting in important changes in key abiotic properties (Crews et al., 1995; Vitousek, 2004; Wardle, Bardgett, et al., 2004). For example, young soils are known to accumulate organic carbon during soil development from centuries to millennia (Schlesinger, 1990), and older soils are expected to support more acid, and P depleted soils compared with younger substrates (Doetterl et al., 2018; Laliberté et al., 2013). Importantly, although soil properties do change as soil develops over geological timescales, the parent material does not vary. Because of this, soil development has been suggested as a good model system to investigate the role of soil abiotic and biotic properties in driving the responses of soil respiration to disturbances such as increasing temperatures (Orwin et al., 2006). A number of studies performed at individual soil chronosequences have investigated whether soil development stage influences soil respiration rates, showing contrasting results. Whereas some studies found an enhancing effect of soil development on soil respiration (J. L. Campbell & Law, 2005; Law, Sun, Campbell, Van Tuyl, & Thornton, 2003), others observed that soil respiration rates decreased as soil develops (Tang et al., 2008; Wang, Bond-Lamberty, & Gower, 2002). These differences are likely due to sitespecific variations in soil development trajectories between chronosequences with contrasting parent material and climatic conditions (Alfaro, Manzano, Marquet, & Gaxiola, 2017). Therefore, to gain a more comprehensive understanding of how soil development affects soil respiration and its response to temperature, such effects should be evaluated both within single chronosequences but also across multiple chronosequences occurring in different ecosystem types with contrasting environmental conditions (e.g. climate, parent material, soil origin, etc.).

Beyond soil properties and soil development, other mechanisms may also modulate soil respiration responses to temperature. For instance, substrate depletion and thermal acclimation have been demonstrated to alter soil respiration responses to temperature (Bradford et al., 2010; Hartley, Hopkins, Garnett, Sommerkorn, & Wookey, 2008). Temperature accelerates microbial activity, leading to an increase in soil respiration (Hochachka & Somero, 2002). However, microorganisms develop several mechanisms to acclimate to the ambient temperature regime such as changes in enzyme and membrane structures. Hence, when subjected to the same temperature range, the microbial activity and soil respiration of acclimated microorganisms would be lower compared to the not acclimated ones (Hochachka & Somero, 2002). Therefore, thermal acclimation to the ambient temperature regime may help to reduce the magnitude of soil respiration responses to temperature (Bradford et al., 2019; Dacal, Bradford, Plaza, Maestre, & García-Palacios, 2019). At the same time, such acceleration in microbial activity with temperature may also cause an important reduction in the availability of readily decomposable C sources, leading to substrate depletion (Cavicchioli et al., 2019; Schindlbacher et al., 2015). Consequently, substrate depletion can limit microbial processes such as soil respiration (Walker et al., 2018). Given that such mechanisms may mitigate soil respiration responses to temperature, they should also be evaluated to improve the accuracy in the predictions of the land C-climate feedbacks.

Herein, we used soil development as an ecological model system to test the importance of soil properties in driving the responses of soil respiration to changes in temperature. To such an end, we take advantage of soils collected from eight chronosequences (Delgado-Baquerizo et al., 2019, 2020) located in Arizona (AZ; USA), California (CAL; USA), Colorado (CO; USA), Hawaii (HA; USA), New Mexico (JOR; USA), Chile (CH), Spain (CI) and Australia (WA) to perform an independent laboratory assay based on short-term soil incubations at three assay temperatures (5, 15 and 25°C). These chronosequences range from hundreds to million years and encompass a wide range of vegetation types (i.e., grasslands, shrublands, and forests), climatic conditions (arid, continental, temperate and tropical), and origins (i.e., sand dunes, sedimentary and volcanic; see Table 1 for more details). Further, we addressed whether soil respiration and its response to temperature change over soil development either within or across chronosequences. Finally, we assessed whether thermal acclimation influences soil respiration responses to temperature across contrasting ecosystem types and soil development stages.

**Table 1.** Climate origin, vegetation type, age, and environmental conditions for eight soil chronosequences. Chronosequence origin describes the major causal agent of each chronosequence. Climate and vegetation types show the main climatic conditions and the dominant vegetation for each chronosequence. MAT= Mean annual temperature, MAP= Mean annual precipitation, SOC= Soil organic carbon, Soil P= Soil phosphorus, and Microbial biomass= Sum of all bacterial, fungi, and other soil microbial biomarkers.

| Chronosequences |  |  |  |  |  |  |  |  |
| --- | --- | --- | --- | --- | --- | --- | --- | --- |
| Label | AZ | CAL | CH | CI | CO | HA | JOR | WA |
| Country | USA | USA | Chile | Spain | USA | USA | USA | Australia |
| Name | SAGA | Merced | Conguillio | La Palma | Coal Creek | Hawaii | Jornada Desert | Jurien Bay |
| Age | 0.9-3000ky | 0.1-3000ky | 0.06-5000ky | 0.5-1700ky | 5-2000ky | 0.3-4100ky | 1.1-25ky | 0.1-2000ky |
| Chronosequence origin | Volcanic | Sedimentary | Volcanic | Volcanic | Sedimentary | Volcanic | Sedimentary | Sand dunes |
| Climate | Arid | Temperate | Temperate | Temperate | Continental | Tropical | Arid | Temperate |
| Vegetation type | Forests | Grasslands | Forests | Forests | Grasslands | Forests | Forblands | Shrublands |
| MAT (°C) | 10.4±1.4 | 16.3±0.3 | 8.7±0.8 | 13.8±1.6 | 9.3±0.5 | 15.9±0.5 | 15.43±0.0 | 19.6±0.1 |
| MAP (mm) | 421±57 | 378±64 | 1907±16 | 451±34 | 482±7 | 1895±380 | 276±4 | 558±4 |
| SOC (%) | 2.6±1.9 | 4.9±2.9 | 3.8±3.5 | 5.1±5.5 | 3.7±1.0 | 25.3±12.5 | 0.6±0.2 | 1.2±0.6 |
| Texture (% clay+silt) | 40.4±28.1 | 44.1±17 | 8.3±2.6 | 23.1±11.7 | 34.6±3.3 | 14.3±3.8 | 18.9±3.5 | 3.8±1.4 |
| pH | 7.2±0.3 | 6±0.8 | 5.8±0.4 | 6.7±0.4 | 6±0.3 | 4.2±0.6 | 8.1±0.4 | 7.3±1.2 |
| Soil P (%) | 0.09±0.02 | 0.06±0.03 | 0.02±0.01 | 0.20±0.05 | 0.06±0.01 | 0.07±0.03 | 0.05±0.01 | 0.02±0.02 |
| Microbial biomass (nmolPLFA/g soil) | 356±371 | 1733±886 | 1293±1752 | 622±738 | 667±289 | 5991±1784 | 126±52 | 112±63 |

## Materials and methods

### Study design and field soil collection

The environmental conditions of the eight chronosequences used spanned a wide gradient in climatic conditions (MAT from 8.7 to 19.55°C, and MAP from 276 to 1907 mm) and soil properties (SOC from 0.6 to 25.3 and the percentage of clay plus silt from 3.8 to 44.1, Table 1). The selected chronosequences included four to six stages of soil development. Stage number one corresponds to the youngest soil, whereas four, five, or six correspond to the oldest one within each chronosequence. Each chronosequence was considered a site, so the total number of sites and stages surveyed in our study is 8 and 41, respectively. At each stage, we established a 50 m x 50 m plot for conducting field surveys. Three parallel transects of 50 m length, spaced 25 m apart, formed the basis of the plot. The total plant cover and the number of perennial plant species (plant diversity) were determined in each transect using the line-intercept method (Delgado-Baquerizo et al., 2019). All of the sites were surveyed between 2016 and 2017 using a standardized sampling protocol (Delgado-Baquerizo et al., 2019). At each plot, three composite soil samples (five soil cores per sample: 0 – 10 cm depth) were collected under the canopy of the dominant ecosystem vegetation type (e.g., grasses, shrubs, and trees). Soil samples were collected during the same days within each soil chronosequence. After field collection, soils were sieved at 2 mm, and a fraction was immediately frozen at −20°C for soil microbial biomass analyses. The rest of the soil was air-dried for a month and used for biochemical analyses and laboratory incubations.

### Soil abiotic properties

We measured the following abiotic soil properties in all samples: soil organic C (SOC), texture (% of clay + silt), pH, and available soil phosphorus (soil P). To avoid confounding effects associated with having multiple laboratories performing soil analyses, all dried soil samples were shipped to Spain (Universidad Rey Juan Carlos) for laboratory analyses. The concentration of SOC was determined by colorimetry after oxidation with a mixture of potassium dichromate and sulfuric acid at 150° C for 30 minutes (Anderson & Ingram, 1993). Soil pH was measured with a pH meter in a 1:2.5 suspensions of dry soil mass to deionized water volume. Soil texture (% clay + silt) was determined on a composite sample per chronosequence stage, according to Kettler, Doran, & Gilbert (2001). Olsen P (soil P hereafter) was determined by extraction with sodium bicarbonate, according to Olsen, Cole, Watanabe, & Dean (1954). Mean annual temperature (MAT) and mean annual precipitation (MAP) values for the soils of each site were obtained using Wordclim version 2.0 (Fick & Hijmans, 2017), which provides global average climatic data for the 1970-2000 period.

### Soil microbial biomass

We estimated soil microbial biomass by measuring phospholipid fatty acids (PLFAs). These were extracted from freeze-dried soil samples using the method described in Bligh & Dyer (1959), as modified by Buyer & Sasser (2012). The extracted PLFAs were analysed on an Agilent Technologies 7890B gas chromatograph with an Agilent DB-5 ms column (Agilent Technologies, CA, USA). The biomarkers selected to indicate total bacterial biomass are the PLFAs i15:0, a15:0, 15:0, i16:0, 16:1ω7, 17:0, i17:0, a17:0, cy17:0, 18:1ω7 and cy19:0, and the biomarker to indicate total fungal biomass is the PLFA 18:2ω6. Using the selected PLFA biomarkers, the biomass was calculated for each soil sample (Frostegård & Bååth, 1996; Rinnan & Bååth, 2009). Total microbial biomass includes the sum of all bacterial and fungal biomarkers plus that of other soil microbial biomarkers such as the eukaryotic C18:1w9.

### Laboratory incubations and soil heterotrophic respiration measurements

We conducted short-term (10 h) incubations of our soil samples, in accordance with previous studies (Atkin & Tjoelker, 2003; Bradford et al., 2010; Hochachka & Somero, 2002; Tucker, Bell, Pendall, & Ogle, 2013), at 5, 15, and 25°C at 60% of WHC. The short timescale used was chosen to prevent acclimation to the assay temperatures used in the laboratory. The incubation temperatures (5, 15 y 25°C) were selected to cover the range spanned by the MAT values of the eight chronosequences studied (from 8.7 to 19.55°C). Additionally, such incubation temperatures are similar to the ones used in previous studies (Bradford et al., 2008, 2019; Dacal et al., 2019). Soil samples were incubated in 96-deepwell microplates (1.3 mL wells) by adding *c.* 0.5 g soil per well. All soil samples were run in triplicate (laboratory replicates). Incubations were performed in growth chambers under dark conditions and 100% air humidity. Microplates were covered with polyethylene film to prevent soil drying but to allow gas exchange.

Soil respiration rates were measured using a modified MicroResp^TM^ technique (C. D. Campbell, Chapman, Cameron, Davidson, & Potts, 2003). Glucose at a dose of 10 mg C g^-1^ dry soil was used as a substrate. It was used to avoid substrate limitation on soil respiration rates (Bradford et al., 2010), as the dose used in our study is supposed to exceed microbial demand (Davidson, Janssens, & Luo, 2006). Soils were incubated at the particular assay temperature (5, 15, and 25°C) for ten hours. However, the detection plates used to measure soil respiration were only incubated during the last 5 hours to avoid the oversaturation of the detection solution. The absorbance of the detection plate was read immediately before and after its use. Three analytical replicates were run per sample, and the mean of these repeats per assay temperature was used as the observation of potential respiration rate for each sample.

### Statistical analyses

We evaluated the importance of soil properties in driving the responses of soil respiration to changes in temperature. To do that, we firstly analysed soil respiration responses to assay temperature within and across chronosequences. For within chronosequences analyses, we built eight linear regression models (LM) including soil development stage, assay temperature, the interaction between both variables, SOC, texture, pH, soil P, and microbial biomass as fixed factors. Soil properties were removed until there is a low collinearity between them and soil development stage (i.e. square-root VIFs <2, Bradford et al., 2017). However, to evaluate the assay temperature effect on soil respiration across chronosequences, we performed a linear mixed-effects model (LMM) with soil development stage (in years), MAT, assay temperature, SOC, texture, pH, soil P, and microbial biomass as fixed factors, and the chronosequence identity as a random factor. We then compared whether there were differences in the magnitude of the effect of assay temperature on soil respiration among chronosequences, using the standardized coefficients of assay temperature obtained in the within chronosequence LMs. Finally, we tested whether biotic and abiotic factors drive the response of soil respiration to temperature. For doing so, we built LMMs that incorporated soil development stage (in years) and assay temperature as fixed factors, and chronosequence identity as a random factor using different subsets of data. Specifically, we grouped the chronosequences in two levels according to each of the environmental conditions and soil properties considered such as the origin of the chronosequence, MAT, SOC, texture, pH, P, and microbial biomass. Then, we ran the model described above separately for each group of data to evaluate how the magnitude of the effect of temperature on soil respiration changes between the models using groups of data with contrasting environmental conditions and soil properties. In most cases, each of the groups of data included four chronosequences each (i.e., half of the chronosequences studied each). We classified each chronosequence by the mean across the whole chronosequence of each of the selected variables to avoid separating different stages of the same chronosequence in different groups. The threshold to distinguish between both groups of each category was established at the value closest to the mean among all observations that allow having the same or almost the same number of chronosequences in each group.

On the other hand, to evaluate the effect of soil development on soil respiration and its response to temperature we used the same approach described above for evaluating the effect of assay temperature on soil respiration (LMs within chronosequences and an LMM across chronosequences). Additionally, we used two different approximations for soil development stage depending on the spatial scale. When analysing each chronosequence separately, we used the stage (from 1 to 6) to address the effects of soil development stage (Delgado-Baquerizo et al., 2019; Laliberté et al., 2013; Wardle, Bardgett, Walker, & Bonner, 2009; Wardle, Walker, & Bardgett, 2004), given the high level of uncertainty in assigning precise ages for many of the chronosequences studied (Wardle, Walker, et al., 2004). However, when analysing across chronosequences, we used the estimation of years as a measure of soil development stage (Crews et al., 1995; Tarlera, Jangid, Ivester, Whitman, & Williams, 2008) to compare chronosequences covering contrasting ranges of soil development stages.

Finally, to test whether the thermal acclimation of soil respiration to the ambient temperature regime influences the soil respiration responses to assay temperature over soil development, we performed an LMM as that described above. We statistically controlled for differences in soil microbial biomass by including it as a covariate in the model (Bradford et al., 2019, 2010; Dacal et al., 2019). All the statistical analyses were conducted using the R 3.3.2 statistical software (R Core Team, 2015). The linear mixed-effects models (LMMs) were fitted with a Gaussian error distribution using the ‘lmer’ function of the lme4 package (Bates, Mächler, Bolker, & Walker, 2015). Response data were transformed by taking the natural logarithm of each value when needed to meet the assumptions of normality and homogeneity of variance.

## Results

### Effects of abiotic and biotic drivers on soil respiration responses to temperature

First, we found a consistent and positive significant effect of assay temperature on soil respiration both within and across chronosequences (P < 0.001 in all cases, Figure 1 and 2, Table S1 and S2, respectively). The magnitude of this positive effect varied between chronosequences (Figure 3). For instance, the assay temperature effect in a Mediterranean sedimentary chronosequence from California (CAL) was 84.5% (95% CI= 51.07%- 117.96%) and 144.44% (95% CI = 94.63% - 146.63%) greater than in a Mediterranean sandy chronosequence in Western Australia (WA) or a volcanic forest chronosequence from Hawaii (HA), respectively (Figure 3).

**Figure 1.**
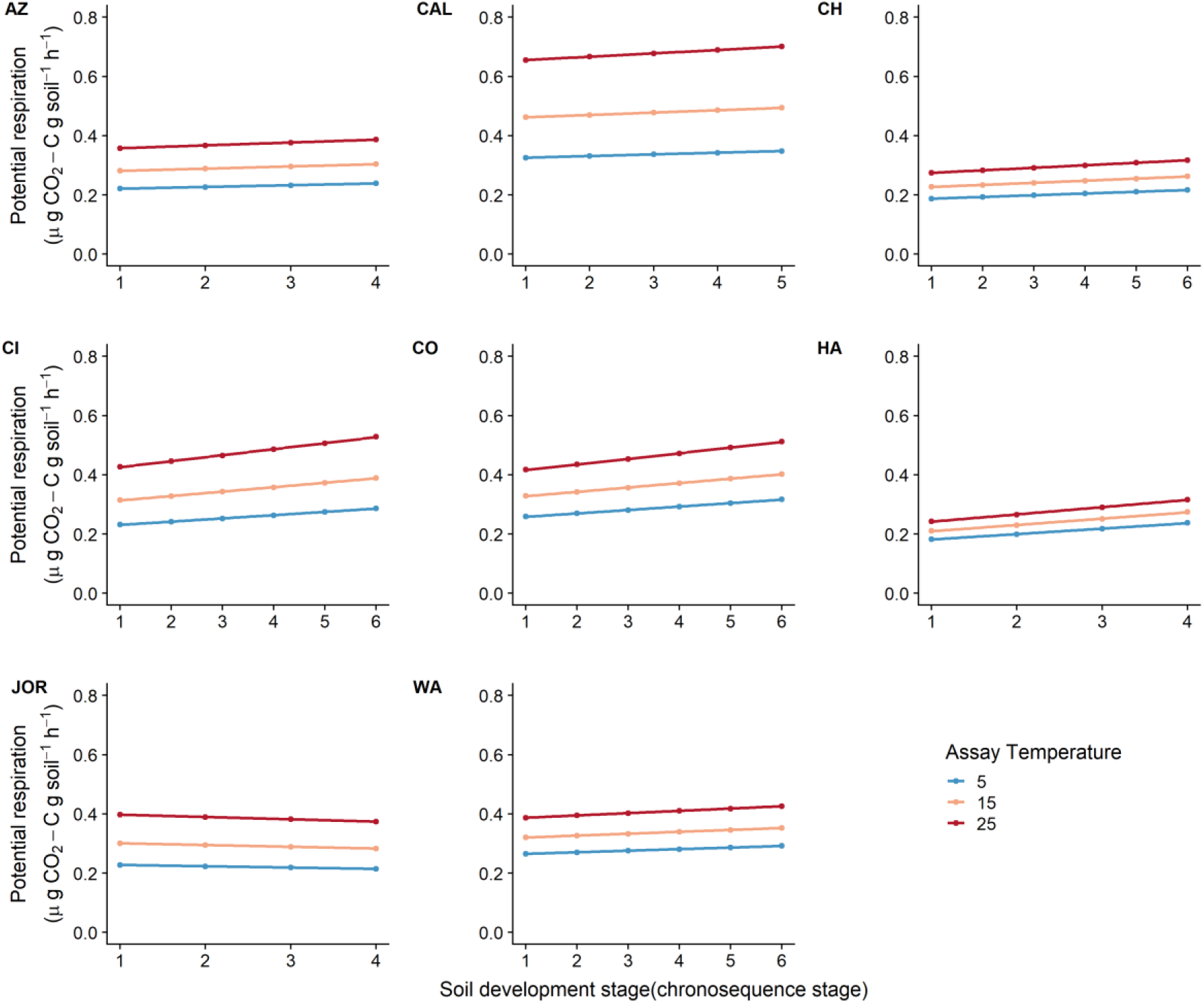
Estimated effects of assay temperature and soil development stage (chronosequence stage) on potential respiration rates at a controlled biomass value and with substrate in excess within chronosequence. The effects were estimated using coefficients from the linear model used for each chronosequence (Table S1). Three outcomes of this model are shown, one for each temperature assayed (i.e. 5, 15, and 25°C). Specifically, we estimated soil respiration rates using the unstandardized coefficients of the model, along with the mean value of the soil properties included in the model of each chronosequence, one of the assay temperatures and one of the soil development stages observed in each chronosequence.

**Figure 2.**
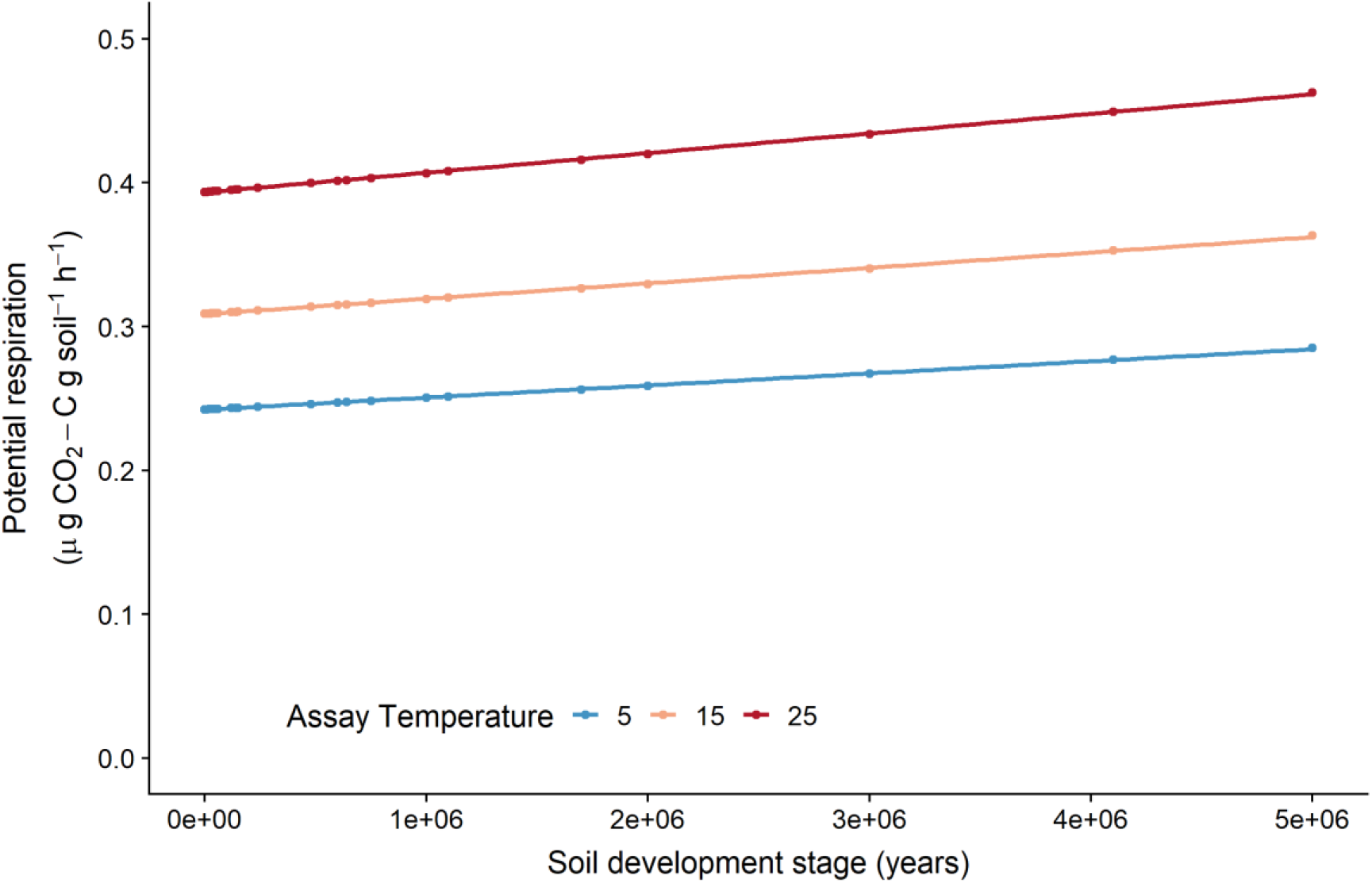
Estimated effects of assay temperature and soil development stage (years) on potential respiration rates at a controlled biomass value and with substrate in excess across chronosequences. The effects were estimated using coefficients from the linear mixed-effects model (Table S2). Three outcomes of this model are shown, one for each temperature assayed (i.e. 5, 15, and 25°C). Specifically, we estimated soil respiration rates using the unstandardized coefficients of the model, along with the mean value of the soil properties included in the model of each chronosequence, one of the assay temperatures and one of the soil development stages observed across all sites.

**Figure 3.**
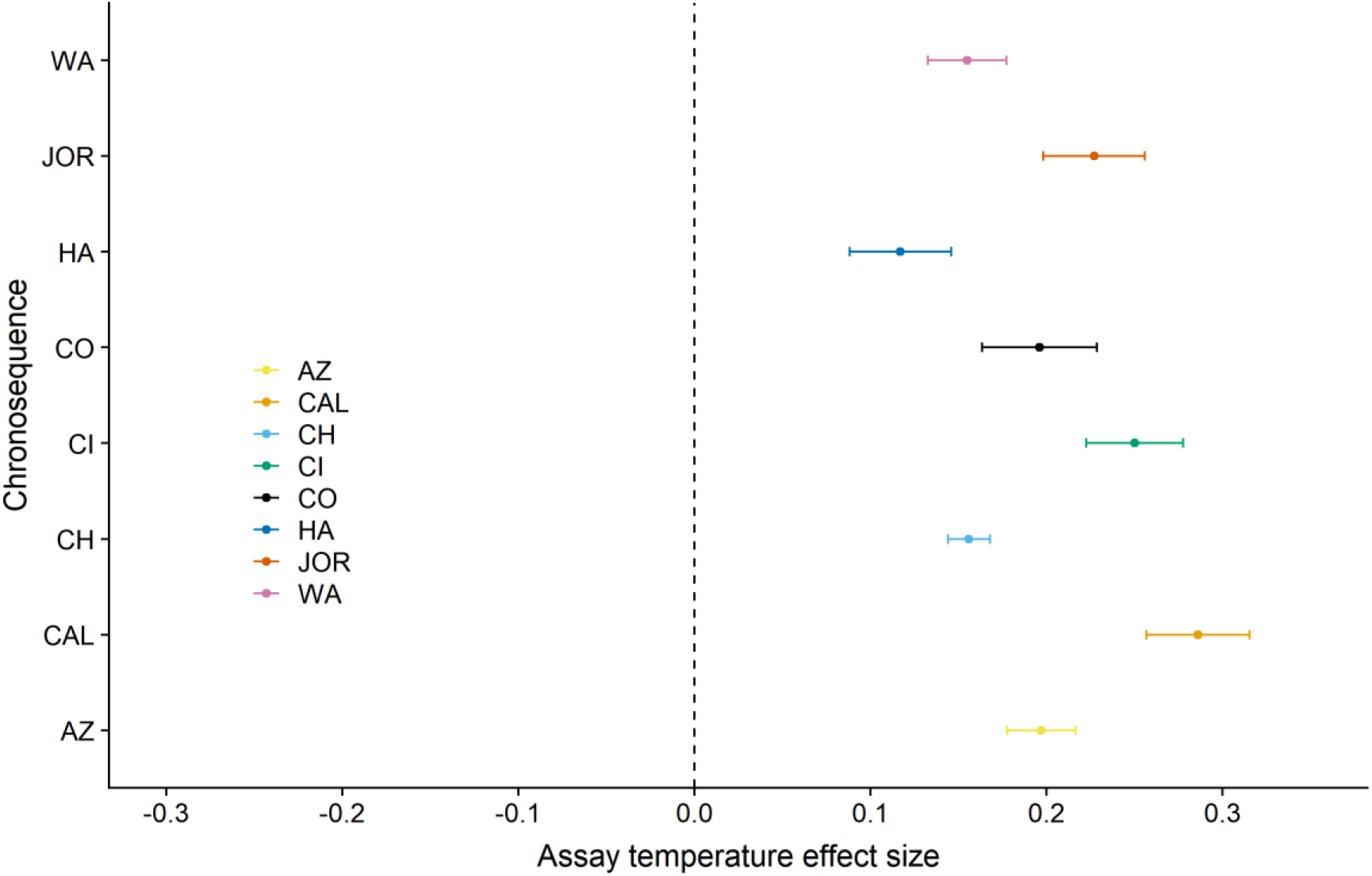
Comparison on the magnitude of the effects of assay temperature on soil respiration among the eight chronosequences studied. The points represent the mean and the error bars correspond to the 95% CI. AZ, JOR, HA presented four stages (n=4), CAL had five stages (n=5) and the rest showed six stages (n= 6).

The effect of assay temperature on soil respiration was consistently positive across all the climatic conditions and soil properties evaluated (Figure 4). However, environmental variables altered the magnitude of the assay temperature effect on soil respiration. For instance, the effect of assay temperature was 12.08% (95% CI = 5.40% - 18.77%) lower for the volcanic chronosequences compared with the ones with a sedimentary or a dune origin (Figure 4). However, the greatest differences on the magnitude of such effect were observed in sites with contrasting soil texture (Figure 4). Specifically, soils with > 20% silt and clay showed a 43.65% (95% CI = 35.18% - 52.12%) higher effect of assay temperature on soil respiration compared with soils with < 20% silt and clay. On the other hand, the effect of assay temperature on soil respiration was 23% (95% CI = 15% - 30%) greater in sites with higher SOC, microbial biomass, and soil P content compared with soils with lower values of such soil properties (Figure 4). The magnitude of the assay temperature effect slight differed (i.e., 9% difference; 95% CI = 5% - 17%) between soils with contrasting pH values (Figure 4). On the other hand, the magnitude of the assay temperature effect on soil respiration did not change across soils with contrasting MAT values (Figure 4).

**Figure 4.**
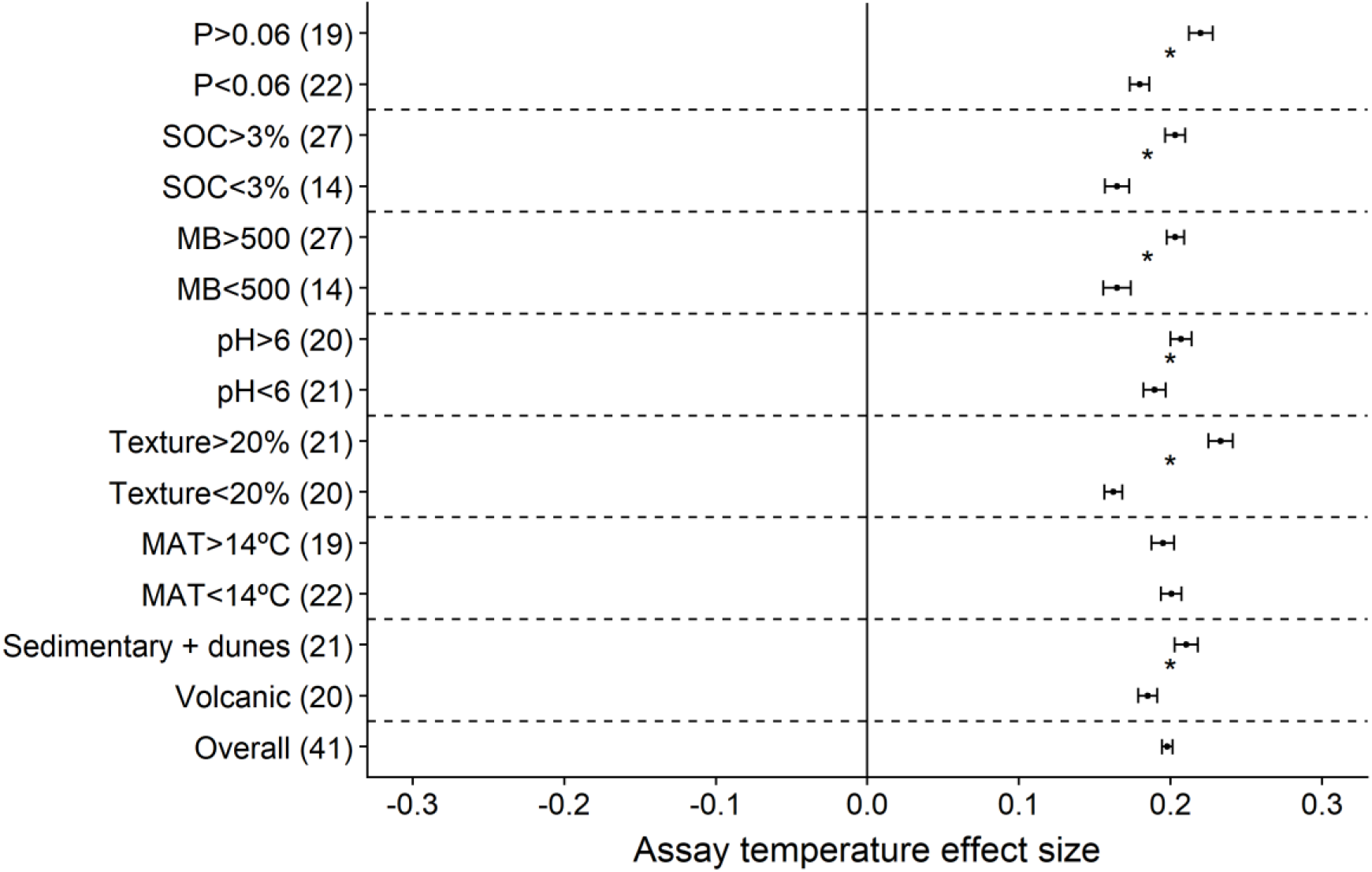
Comparison of the effects of assay temperature on soil respiration among different environmental conditions. The points represent the mean and the error bars correspond to the 95% CI. Asterisks denote significant differences at *p* < 0.05. The total n was shown in brackets and it was the result of the number of stages within the chronosequences x the number of chronosequences included in each level of the classification. MAT= mean annual temperature, Texture=% of clay + silt, MB= total microbial biomass, SOC= soil organic carbon, and P= soil phosphorus. Volcanic and sedimentary + dunes refer to the different origins observed across the eight chronosequences studied.

### Effect of soil development on soil respiration and its response to temperature

When analysing the effect of soil development on soil respiration at every chronosequence separately, we did not observe any significant effect in five out of eight chronosequences (Figure 1, Table S1). We found higher soil respiration rates in older soils than in younger ones in three volcanic chronosequences located in temperate and tropical forests in Chile (i.e., CH, P = 0.016, Figure 1, Table S1), Spain (i.e., CI, P = 0.049, Figure 1, Table S1) and Hawaii (i.e., HA, P = 0.009, Figure 1, Table S1). We also observed a positive effect of soil development on respiration across chronosequences (P = 0.004, Figure 2, Table S2). Regardless these results, soil development did not affect respiration responses to temperature neither within nor across chronosequences, as the interaction between soil development and assay temperature was not significant (P > 0.05 in all cases).

### Thermal acclimation of soil respiration to ambient temperature regimes

The site MAT did not affect soil respiration (P = 0.487, Table S2) nor its response to assay temperature (MAT × assay temperature, P = 0.807), suggesting the absence of acclimation of soil respiration to the ambient temperature regime. The lack of MAT effect on soil respiration was constant across all soil development stages (MAT × soil development, P = 0.122).

## Discussion

Our study shows that elevated temperatures consistently increased soil heterotrophic respiration rates across contrasting soil chronosequences. Although older soils tended to support higher soil respiration–especially in volcanic, temperate, and tropical forests–, our findings indicate that soil development did not alter the relationship between heterotrophic respiration and temperature. Conversely, soil properties such as SOC, the amount of clay and silt, pH, microbial biomass, and P content had a significant control on the magnitude of positive temperature effects on soil respiration. Overall, these findings provide new insights into the role of soil properties in driving soil respiration responses to temperature, which are essential to project the magnitude of the land C-climate feedback accurately.

We observed a consistent positive effect of assay temperature on soil respiration within and across chronosequences. Such results agree with previous literature addressing the effects of temperature on soil organic matter decomposition and soil respiration rates (Davidson & Janssens, 2006; Kirschbaum, 2006; Lloyd & Taylor, 1994; Min et al., 2020). The enhancing effect of temperature on soil respiration is largely driven by the acceleration of microbial metabolic rates (Hochachka & Somero, 2002). Importantly, the effect of elevated temperatures on soil respiration was positive in all chronosequences studied, suggesting that this enhancing effect, at least in our study, is independent of the ecosystem type. However, certain chronosequences showed differences in the magnitude of the assay temperature effect between them. That could be explained by our results indicating that environmental conditions and soil biotic and abiotic properties have the ability to determine the magnitude of the consistently positive effect of temperature on soil respiration. For instance, soil respiration responses to assay temperature differed depending on the origin of the chronosequence considered. Such results suggest that parent material also influences soil respiration responses to temperature. An explanation for these observed differences could be that soil develops differently according to several factors such as soil parent material (Alfaro et al., 2017; Carlson, Flagstad, Gillet, & Mitchell, 2010; Jenny, 1941). Moreover, we found that the magnitude of the effect of assay temperature was lower in sites with less soil P available. Such results indicate that this nutrient is necessary to sustain microbial activity (Liu, Gundersen, Zhang, & Mo, 2012). Further, we also observed differences in the magnitude of the response of soil respiration to elevated temperatures between sites with contrasting amounts of clay and silt. These differences could be caused by the fact that water availability in the soil is expected to increase when the amount of clay and silt in the soil rises (Delgado-Baquerizo et al., 2013), accelerating microbial activity (Karhu et al., 2014; Luo, Wan, Hui, & Wallace, 2001). However, this effect of the amount of clay and silt on soil respiration responses to temperature could disappear at high amounts of clay and silt, as clay and silt may limit microbial access to SOC. Also, the magnitude of the effect of assay temperature on soil respiration increased in sites with greater soil pH, as the microbial activity is negatively affected by acidification (Reth, Reichstein, & Falge, 2005; Rustad et al., 2000). Finally, our results indicated that soil respiration response to assay temperature increases with substrate availability (i.e., SOC) and microbial biomass. This increase in soil respiration rates in response to temperature under high SOC and microbial biomass conditions may cause the acceleration of microbial activity and, subsequently, a substrate depletion and an important reduction of microbial biomass (Cavicchioli et al., 2019). Thus, our findings provide new insights about how soil properties modulate the magnitude of the consistently enhancing effect of temperature on soil respiration.

In three out of the eight chronosequences evaluated, we found a significant positive effect of soil development on soil respiration rates. Interestingly, all these chronosequences shared a volcanic origin. The different effect of soil development on soil respiration found across chronosequences may be mediated by contrasting parent material between them, leading to variations in the soil development trajectories followed by the eight chronosequences evaluated. The differences in the range of years covered by each of the chronosequences evaluated may also influence the effect of soil development on soil respiration. Such contrasting results observed when analysing each chronosequence separately limits our capacity to draw more general conclusions about how soil C losses to the atmosphere via soil respiration change over soil development, specially under a warming scenario. Such limitations are similar to the ones found in previous studies (J. L. Campbell & Law, 2005; Law et al., 2003; Saiz et al., 2006; Tang et al., 2008; Wang et al., 2002) conducted on a single chronosequence and covering a narrow range of soil development stages (from years to centuries). Therefore, when evaluating soil development effect on soil respiration across chronosequences, we observed a significant enhancing effect of soil development stage on soil respiration. Our findings improve our knowledge about the effect of soil development stage on soil respiration across large spatial scales including different ecosystem types with contrasting environmental conditions and soil properties. Specifically, our results indicated that elder soils have greater soil C losses to the atmosphere than younger ones. Such greater soil respiration rates found in elder soils within some and across chronosequences may be explained by the increase in soil C easily releasable from mineral-SOC associations in soils that had experienced higher weathering (Keiluweit et al., 2015). Conversely, we observed that soil development did not modulate the magnitude of the effect of assay temperature on soil respiration, as the interaction between soil development stage and assay temperature was not significant either within or across chronosequences. These results indicate that, no matter how old soils are, soil carbon stocks are highly sensitive to increases in temperature associated with climate change. Thus, although worldwide soils show contrasting ages (Laliberté et al., 2013; Wardle, Bardgett, Walker, Peltzer, & Lagerström, 2008), they present similar soil respiration responses to temperature. Further, the assay temperature effect was at least three times larger in magnitude than the effect of soil development stage on soil respiration. Such results agree with previous studies showing pronounced soil respiration responses to assay temperature (Bradford et al., 2010), especially across large temperature ranges such as those used in our incubations (i.e. from 5 to 25°C). Consequently, our study supports that soil microbial communities from very different ecosystem types are capable of rapidly responding to increasing temperature, resulting in greater soil respiration.

A growing body of evidence suggests that thermal acclimation of soil microbial respiration to temperature can be found across large spatial scales (Bradford et al., 2019, 2010; Dacal et al., 2019; Ye, Bradford, Maestre, Li, & García-Palacios, 2020). However, we did not find a significant effect of MAT, suggesting that soil respiration is not acclimated to the ambient temperature regime at our sites. This apparent disagreement may be due to the shorter MAT gradient evaluated in our study (i.e., from 8.7°C to 19.55°C) compared with previous ones (i.e., from −2 to 28°C; Bradford et al., 2019; Dacal et al., 2019; Ye, Bradford, Maestre, Li, & García-Palacios, 2020). Nevertheless, our results are similar to other cross-biome studies (Carey et al., 2016; Karhu et al., 2014), and may be the result of negligible effects of thermal acclimation on soil respiration when compared with overarching factors such as assay temperature (Hochachka & Somero, 2002).

In conclusion, we found that assay temperature consistently enhanced soil respiration across contrasting chronosequences. On the other hand, we observed no evidence of thermal acclimation of soil respiration to the ambient temperature regime. Although we observed a positive effect of soil development on soil respiration, it did not change the magnitude of the assay temperature effect. Despite the clear and positive effect of assay temperature on soil respiration observed, soil properties such as SOC, texture, pH, P content, and microbial biomass significantly modified the magnitude of this positive soil respiration response to temperature. Our findings emphasize the role of biotic and biotic soil properties as drivers of soil respiration responses to temperature across biomes and provide new insights to better understand the magnitude of the land C-Climate feedback and to establish accurate greenhouse emission targets.

## Acknowledgements

This project received funding from the European Union’s Horizon 2020 research and innovation program under Marie Sklodowska-Curie Grant Agreement 702057. M.D. was supported by an FPU fellowship from the Spanish Ministry of Education, Culture and Sports (FPU-15/00392). M.D. and F.T.M. are supported by the European Research Council (Consolidator Grant Agreement No 647038, BIODESERT). M.D-B. is supported by a Large Research Grant from the British Ecological Society (grant agreement n° LRA17\1193, MUSGONET). F.T.M and M.D-B. acknowledge support from the Spanish Ministry (project CGL2017-88124-R). PGP and M.D-B. are supported by a Ramón y Cajal grant from the Spanish Ministry of Science and Innovation (RYC2018-024766-I and RYC2018-025483-I, respectively). F.T.M. acknowledges support from the Generalitat Valenciana (CIDEGENT/2018/041). We would like to thank Matt Gebert, Jessica Henley, Victoria Ochoa, and Beatriz Gozalo for their help with lab analyses. We also want to thank Lynn Riedel, Julie Larson, Katy Waechter and Drs. David Buckner and Brian Anacker for their help with soil sampling in the chronosequence from Colorado, and to the City of Boulder Open Space and Mountain Parks for allowing us to conduct these collections.

## Authorship

M.D., M.D.-B. and P.G.P developed the original idea of the analyses presented in the manuscript. M.D.-B. designed the field study and wrote the grant that funded the work. J.B. conducted the laboratory work with inputs from M.D.-B and A.G. M.D. performed the statistical analyses, with inputs from M.D.-B., F.T.M and P.G.P. All authors included A.A.B. contributed to data interpretation. M.D. wrote the first version of the manuscript, which was revised by all co-authors.

## Competing interests

The authors declare no competing financial interests.

## Notes

### Competing Interest Statement

The authors have declared no competing interest.

