## Supplementary material for "Temperature increases soil respiration across ecosystem types and soil development, but soil properties determine the magnitude of this effect": Table S

**Table S1.** Summary results of the linear regression models (LMs) used to test the effect of temperature and the chronosequence stage on soil respiration within each of the eight chronosequences studied. The table shows the standardized coefficients (mean ± SD) of the model. Respiration rates were ln-transformed to meet normality assumptions. AZ, JOR, HA presented four stages (total n=4), CAL had five stages (total n=5) and the rest showed six stages (total n= 6). P values below 0.05 are shown in bold. The coefficients of the table were used to create Figure 1 and Figure 3.

| Chronosequence | Variable | Coefficients | t value | P value | $R^2$ |
|---|---|---|---|---|---|
| **AZ** | Intercept | -0.946 ± 0.111 | -8.58 | **<0.001** | 0.78 |
| | Assay temperature | 0.197 ± 0.020 | 9.91 | **<0.001** | |
| | Stage | 0.041 ± 0.032 | 1.27 | 0.214 | |
| | SOC | 0.264 ± 0.119 | 2.22 | **0.035** | |
| | pH | -0.128 ± 0.109 | -1.17 | 0.252 | |
| | Total biomass | 0.195 ± 0.136 | 1.43 | 0.164 | |
| **CAL** | Intercept | -0.756 ± 0.063 | -12.07 | **<0.001** | 0.65 |
| | Assay temperature | 0.286 ± 0.034 | 8.21 | **<0.001** | |
| | Stage | 0.027 ± 0.061 | 0.438 | 0.664 | |
| | Texture | -0.046 ± 0.060 | -0.76 | 0.452 | |
| | Total biomass | 0.283 ± 0.124 | 2.27 | **0.029** | |
| **CH** | Intercept | -1.357 ± 0.094 | -14.47 | **<0.001** | 0.71 |
| | Assay temperature | 0.156 ± 0.015 | 10.42 | **<0.001** | |
| | Stage | 0.046 ± 0.018 | 2.51 | **0.016** | |
| | SOC | 0.086 ± 0.047 | 1.82 | 0.075 | |
| | Texture | 0.060 ± 0.125 | 0.48 | 0.631 | |
| **CI** | Intercept | -1.108 ± 0.101 | -10.96 | **<0.001** | 0.48 |
| | Assay temperature | 0.25 ± 0.034 | 7.28 | **<0.001** | |
| | Stage | 0.068 ± 0.033 | 2.02 | **0.049** | |
| | pH | 0.093 ± 0.102 | 0.908 | 0.368 | |
| | Soil P | 0.010 ± 0.046 | 0.228 | 0.821 | |
| **CO** | Intercept | -0.907 ± 0.131 | -6.93 | **<0.001** | 0.3 |
| | Assay temperature | 0.196 ± 0.041 | 4.79 | **<0.001** | |
| | Stage | 0.064 ± 0.043 | 1.50 | 0.14 | |
| | pH | 0.165 ± 0.198 | 0.83 | 0.41 | |
| | Soil P | -0.344± 0.383 | -0.90 | 0.374 | |
| | Total biomass | 0.452 ± 0.354 | 1.28 | 0.207 | |
| **HA** | Intercept | -1.302 0.137 | -9.52 | **<0.001** | 0.38 |
| | Assay temperature | 0.117 ± 0.03 | 3.98 | **<0.001** | |
| | Stage | 0.140 ± 0.05 | 2.78 | **0.009** | |
| | Texture | -0.069 ± 0.183 | -0.38 | 0.707 | |
| | Total biomass | -0.039 ± 0.035 | -1.12 | 0.27 | |
| **JOR** | Intercept | 0.329 ± 0.921 | 0.357 | 0.724 | 0.64 |
| | Assay temperature | 0.227 ± 0.03 | 7.71 | **<0.001** | |
| | Stage | -0.032 ± 0.043 | -0.74 | 0.465 | |

| | | | | | |
|---|---|---|---|---|---|
| | SOC | 1.803 ± 1.363 | 1.32 | 0.196 | |
| | Texture | 0.335 ± 0.162 | 2.07 | **0.048** | |
| | Total biomass | 0.802 ± 1.145 | 0.70 | 0.489 | |
| **WA** | Intercept | -1.117 ± 0.524 | -2.13 | **0.038** | 0.35 |
| | Assay temperature | 0.155 ± 0.028 | 5.51 | **<0.001** | |
| | Stage | 0.030 ± 0.027 | 1.13 | 0.265 | |
| | Texture | -0.307 ± 0.465 | -0.66 | 0.512 | |
| | Total biomass | 0.510 ± 1.103 | 0.661 | 0.512 | |

**Table S2.** Summary results of the linear mixed-effect regression model (LMM) to test the effect of temperature and soil age on soil respiration across all chronosequences. The table shows the unstandardized coefficients (mean ± SD) of the model. Respiration rates were ln-transformed to meet normality assumptions. We used three replicates per chronosequence stage, eight chronosequences (with 4, 5 or 6 stages) and three assay temperatures (total n = 364). P values below 0.05 are shown in bold. The coefficients of the table were used to create Figure 2.

| Variable | Coefficients | df | t value | P value |
|---|---|---|---|---|
| **Intercept** | -1.155±0.082 | 6 | -14.022 | **<0.001** |
| **Assay temperature** | 0.198±0.012 | 349 | 17.142 | **<0.001** |
| **MAT** | 0.064±0.086 | 6 | 0.743 | 0.487 |
| **Soil age** | 0.039±0.013 | 351 | 2.930 | **0.004** |
| **SOC** | 0.022±0.027 | 353 | 0.828 | 0.408 |
| **Texture** | 0.035±0.019 | 351 | 1.799 | 0.073 |
| **pH** | 0.020±0.026 | 349 | 0.773 | 0.440 |
| **Soil P** | 0.015±0.029 | 284 | 0.514 | 0.608 |
| **Total biomass** | 0.011±0.029 | 355 | 0.387 | 0.699 |
